# Cardiac fibroblasts regulate cardiomyocyte hypertrophy through dynamic regulation of type I collagen

**DOI:** 10.1101/2022.05.25.493406

**Authors:** Qinghang Meng, Stephanie L. K. Bowers, Yasuhide Kuwabara, Jiuzhou Huo, Rachel Minerath, Allen J. York, Michelle A. Sargent, Vikram Prasad, Anthony J. Saviola, David Ceja Galindo, Kirk C. Hansen, Ronald J. Vagnozzi, Katherine E. Yutzey, Jeffery D. Molkentin

**Author notes:** To whom correspondence should be addressed., Cincinnati Children’s Hospital Medical Center, Howard Hughes Medical Institute, Molecular Cardiovascular Biology, 240 Albert Sabin Way, MLC 7020, Cincinnati, OH 45229 USA. These authors contributed equally (Qinghang Meng, Stephanie L. K. Bowers).

## Abstract

**Rationale:** Cardiomyocytes and fibroblasts in the heart communicate through both secreted growth factors as well as through sensing the structural properties of the extracellular matrix that each helps generate. Previous studies have shown that defects in fibroblast activity during disease stimulation result in altered cardiomyocyte hypertrophy, although the role that collagen might play in this communication is unknown.

**Objective:** Here we investigated how type I collagen maturation and disease-responsive matrix expansion in the heart by cardiac fibroblasts impacts cardiac fibrosis and cardiomyocyte hypertrophy.

**Methods and Results:** We generated and characterized *Col1a2*^*-/-*^ mice using standard gene-targeting. *Col1a2*^*-/-*^ mice were viable, although by young adulthood their hearts showed alterations in extracellular matrix mechanical properties, as well as an unanticipated activation of cardiac fibroblasts and induction of a progressive fibrotic response. This included increases in fibroblast number and a progressive cardiac hypertrophy, with reduced functional performance by 9 months. *Col1a2-loxP* targeted mice were also generated and crossed with the tamoxifen-inducible *Postn-MerCreMer* knock-in mice to delete the *Col1a2* gene in myofibroblasts post-pressure overload injury, to more specifically implicate fibroblasts as effectors of cardiomyocyte hypertrophy in vivo. Opposite to the gradual induction of cardiac hypertrophy observed in germline *Col1a2*^-/-^ mice as they matured developmentally, adult fibroblast-specific deletion of *Col1a2* during pressure overload protected these mice from cardiac hypertrophy in the first week with a delayed fibrotic response. However, this reduction in hypertrophy due to myofibroblast-specific *Col1a2* deletion was gradually lost over 2 and 6 weeks of pressure overload as augmented fibrosis returned.

**Conclusions:** Defective type I collagen in the developing heart alters the structural integrity of the extracellular matrix that leads to fibroblast expansion, activation, fibrosis and hypertrophy with progressive cardiomyopathy in adulthood. However, acute deletion of type I collagen production for the first time in the adult heart during pressure overload prevents ECM expansion and inhibits cardiomyocyte hypertrophy, while gradual restoration of fibrosis again permitted hypertrophy comparable to controls.

## Introduction

The extracellular matrix (ECM) is a highly dynamic noncellular structure that surrounds cells in essentially all vertebrate solid and soft tissues. Excessive ECM deposition in the heart, more generally referred to as cardiac fibrosis, is a common feature of cardiac disease.^1^ To elucidate the underlying mechanisms regulating cardiac fibrosis, studies have focused on dissecting the contribution of individual cell types in ECM protein production. Indeed, the cardiac fibroblast has been implicated as the primary cell type regulating the ECM compartment of the heart, and more specifically the content and quality of the collagen networks. With acute injury or chronic disease states of the heart fibroblasts convert to a more highly differentiated and contractile cell known as a myofibroblast, which more efficiently expands the ECM network and total collagen content.^2^ Indeed, defective collagen production by cardiac fibroblasts due to deletion of either heat shock protein 47 (HSP47)^3^ or periostin^4^ renders the heart unable to induce an effective fibrotic response, which secondarily impacts the ability of cardiomyocytes to hypertrophy with disease stimuli. Similarly, disrupting the activity of TGFβ receptors 1 and 2 in cardiac fibroblasts, which inhibited the disease-induced fibrotic response, reduced the ability of cardiomyocytes in the heart to hypertrophy.^5, 6^ These previous results suggest that activated fibroblasts are important in mediating cardiac hypertrophy in conjunction with their ability to augment ECM content.

The ECM in the adult heart is primarily composed of type I collagen.^7^ The triple-helix, rod-like type I collagen is formed by two α1(I) and one α2(I) procollagen chains encoded by the *Col1a1* and *Col1a2* genes, respectively.^8, 9^ The osteogenesis imperfecta mutant mouse (OIM), which has a recessive mutation in the *Col1a2* gene, models the human connective tissue disorder osteogenesis imperfecta. Due to the mutation, the OIM/OIM homozygous mice have a severe reduction in functional collagen 1a2.^10^ As a compensatory response, collagen 1a1 chains can homotrimerize, although this generates collagen that is substantially weaker.^10, 11^ In the heart, collagen fibers from OIM/OIM mice were smaller in diameter and less dense.^12^ The left ventricles of these mice were more compliant and readily expanded with passive inflation of a balloon.^12^ Hence, the homotrimeric collagen 1a1 fibrils that characterizes the OIM mouse, while sufficient to support viability, are nonetheless structurally defective.^11^ Indeed, the OIM/OIM mice have a high mortality rate following myocardial infarction (MI) due to rupturing of the myocardial wall.^13^

While these studies showed the importance of type I collagen as a structural component of the ECM in the heart, they did not investigate how such alterations impacted communication between cardiomyocytes and fibroblasts in response to disease stimuli. Here we investigated how fibroblast-generated type I collagen underlies structural rigor of the ECM to protect against baseline disease, or how the ability to quickly expand type I collagen with disease permits effective cardiac hypertrophy. We determined that developmental loss of a structurally rigorous type I collagen-containing ECM network in the heart leads to a secondary reactive expansion of cardiac fibroblasts, progressive fibrosis with periostin induction, and cardiomyopathy with loss of cardiac function over time. More intriguingly, acute deletion of *Col1a2* from pressure overload-activated cardiac fibroblasts prevented new collagen production over 1 weeks’ time that was associated with an inhibition of the hypertrophic response, while gradual production of defective collagen 1a1 homotrimers eventually compensated and hypertrophic potential was restored. Thus, the quality and quantity of type I collagen in the heart underlies structural ECM properties that cardiomyocytes directly sense, both at baseline and in response to disease stimuli, which then impacts their hypertrophic potential.

## Material and Methods

### Animals and surgical models

Mouse founders carrying the *Col1a2* knockout first allele (*Col1a2*^*-/-*^) were purchased from Mutant Mouse Resource & Research Centers (MMRRC, ID: 037695-UCD). To generate *Col1a2 lox-P* only targeted mice, the *Col1a2*^*-/-*^ mouse was further crossed with *Rosa26-Flpe* females (B6.129S4-*Gt(ROSA)26Sor*^*tm1(FLP1)Dym*^/RainJ) to remove the neomycin cassette at the frt sites as described previously.^14^ The *Tcf21-MerCreMer* (MCM) and *Postn-MerCreMer* mice have been previously described respectively.^14, 15^ The *Rosa26 loxP* site–dependent reporter mice (*R26*^*NG-loxP-eGFP*^) were purchased from the Jackson Laboratories (stock no. 012429).^16^ All crosses were carried out and maintained in the C57/BL6 background unless otherwise noted. To induce Cre activity in *Tcf21*^*MCM*^ or *Postn*^*MCM*^ mice *in vivo*, mice were fed tamoxifen-citrate chow (40 mg/kg body weight, Envigo-TD.130860) or were treated with tamoxifen (Millipore Sigma, T5648) that was dissolved in corn oil through i.p. injections (75 mg/kg body weight/day). Transverse aortic constriction (TAC) surgery was performed to induce pressure overload and cardiac hypertrophy as described previously.^17^ Briefly, a silk ligature was tied around a 26-gauge wire and the mouse transverse aorta to produce a pressure load on the heart. Myocardial infarction (MI) was induced by permanent ligating the left coronary artery as described previously.^18^ Animals were handled in accordance with the principles and procedures of the Guide for the Care and Use of Laboratory Animals. All proposed procedures were approved by the Institutional Animal Care and Use Committee at Cincinnati Children’s Hospital. Animal groups and experiments were handled in a blinded manner where possible, and randomization was not needed given that the selected genotypes were all otherwise genetically identical and given that the same ages and equal ratio of sexes were used.

### Antibodies and related reagents

Antibodies against the following proteins were used: periostin (Novus Biologicals NBP1-30042; 1:300 dilution for IF, 1:1000 for Western blot); collagen I (Abcam ab21286; 1:100 for IF); PDGFRα from (R&D Systems AF1062; 1:1000 for IF); collagen 1a2 (Santa Cruz sc-393573; 1:500 for Western blot) Anti-CD31 was from BioLegend (102423; 1:100 for flow cytometry); anti-CD45 was from BD Biosciences (563890; 1:100 for flow cytometry); anti-MEFSK4 was from Miltenyi Biotec (130-120-802; used 1:30 for flow cytometry). Wheat germ agglutinin (488) was from ThermoFisher (W11261; 5 µg/ml for IF).

### Isolation of cardiac fibroblasts

Whole hearts isolated from anesthetized mice were briefly rinsed in cold sterile 1X PBS and the atria were removed. Whole ventricles were minced on ice using surgical scissors into approximately 2 mm pieces (8-10 pieces per mouse heart). Each dissociated ventricle was digested in 2 mL of DMEM containing 2 mg/ml Worthington collagenase type IV (#LS004188), 1.2 U/ml dispase II (Roche, #10165859001), 0.9 mM CaCl^2^ and 2% fetal bovine serum (FBS) at 37°C for 20 min with gentle rotation followed by manual trituration 12-15 times with a 10 mL serological pipette, such that all the tissue pieces were able to pass through the pipette. The tissues were then settled by sedimentation and the supernatant was passed through a 40 µm mesh strainer and stored on ice. Two milliliters of fresh digestion buffer were added, followed by 2 additional rounds of incubation, trituration and replacement of supernatant with fresh digestion buffer, except trituration was performed with a 5 ml serological pipette for round 2 and a 1 ml p1000 pipette tip (USA Scientific, #1112-1720) for round 3. The pooled supernatant from the 3 rounds of digestion was washed with sterile 1X PBS (add 1X PBS to make a final volume of 40 ml) and centrifuged at 200 Xg for 20 min at 4 °C in a swinging bucket rotor centrifuge without brakes. The pellet was resuspended in Red Blood Cell Lysis buffer (155 mM NH_4_Cl; 12 mM NaHCO_3_; 0.1 mM EDTA) for 5 min at room temperature and centrifuged at 200 Xg for 20 min at 4 °C in a swinging bucket rotor centrifuge without brakes. The pellet was then resuspended in flow cytometry sorting buffer consisting of 1X HBSS supplemented with 2% bovine growth serum (BGS) and 2 mM EDTA.

### Flow Cytometry

Dissociated heart suspensions were prepared as described immediately above, and the cells were labeled with antibodies as also described above and were then analyzed using a BD FACSCanto cytometer running BD FACSDiVa software (BD Biosciences), using a violet (405 nm) laser to detect the Brilliant Violet 421-conjugated CD31 and CD45 antibodies and a red (633 nm) laser to detect the APC-conjugated MEFSK4 antibody (antibody concentrations given above). Initial gating with forward and side scatter was used to define single cells. Analysis and quantitation were defined as the total cell number of cells per milligram of dissociated tissue and was performed using FlowJo software from Tree Star, Inc.

### RNA extraction and Western blots

Whole ventricles or isolated cells were digested in Trizol (Thermo Fisher Scientific, Cat. #15596018) for mRNA isolation. cDNA was synthesized using iScript cDNA synthesis kit (Bio-Rad) following the manufacturer’s protocol. Whole ventricles or isolated cells were digested in RIPA buffer with cOmplete™ Mini protease inhibitor cocktail (Millipore Sigma, Cat.# 4693124001) and PhosSTOP phosphatase inhibitor cocktail (Roche Diagnostics, Cat.# 4906837001) for protein isolation. Ventricle ECM proteins were isolated using Subcellular Protein Fractionation Kit for Tissues (Thermo Fisher Scientific, Cat. #87790) following the manufacture’s instruction. Equal amounts of protein were used for SDS gel electrophoresis and Western blots.

### RNA microarray

Whole mouse heart ventricles collected from either *Col1a2*^*+/-*^ or *Col1a2*^*-/-*^ at 2 months of age were lysed in Trizol for total RNA isolation followed by RNA quality assessed using an Agilent 2100 Bioanalyzer. Microarray analysis was performed using the Affymetrix Clariom S platform at Cincinnati Children’s Hospital Medical Center Gene Expression Core Facility. Data in CHP files was analyzed using Transcriptome Analysis Console (Applied Biosystems), Clariom_S_Mouse TAC Configuration file, and iPathwayGuide (Advaita Bioinformatics) to determine differential gene expression between experimental groups. The raw RNA microarray expression data were submitted to the GEO omnibus with an accession number of GSE204724 (embargoed until accepted for publication at mainstream journal)

### ECM mass spectrometry

Preparation of tissue samples was performed as previously described.^19^ Briefly, 5 mg of lyophilized tissue samples were processed by a stepwise extraction with CHAPS and high salt, guanidine hydrochloride and chemical digestion with hydroxylamine hydrochloride (HA) in Gnd-HCl generating cellular, soluble ECM (sECM), and insoluble ECM (iECM) fractions for each sample, respectively. Protein concentration of each fraction for each sample was measured using A660 Protein Assay (Pierce™). Thirty micrograms of protein resulting from each fraction was subjected to proteolytic digestion using a filter-aided sample preparation (FASP) protocol ^20^ with 10 kDa molecular weight cutoff filters (Sartorius Vivacon 500 #VN01H02). Samples were reduced with 5 mM tris(2-carboxyethylphosphine), alkylated with 50 mM 2-chloroacetamide, and digested overnight with trypsin (enzyme:substrate ratio 1:100) at 37ºC. Peptides were recovered from the filter using successive washes with 0.2% formic acid (FA). Aliquots containing 10 μg of digested peptides were cleaned using PierceTM C18 Spin Tips (Thermo Scientific, Cat. #84850) according to the manufacturer’s protocol, dried in a vacuum centrifuge, and resuspended in 0.1% formic acid in mass spectrometry-grade water.

Liquid chromatography mass spectrometry (LC-MS/MS) was performed using an Easy nLC 1200 instrument coupled to a Orbitrap Fusion Lumos Tribrid mass spectrometer (all from ThermoFisher Scientific) as previously described.^19^ Fragmentation spectra were searched against the UniProt *Mus musculus* proteome database (Proteome ID # UP000000589 downloaded 1 December 2021) using the MSFragger-based FragPipe computational platform.^21^ Contaminants and reverse decoys and were added to the database automatically. The precursor-ion mass tolerance and fragment-ion mass tolerance were set to 10 ppm and .2 Da, respectively. Fixed modifications were set as carbamidomethyl (C), and variable modifications were set as oxidation (M), oxidation (P) (hydroxyproline), Gln->pyro-Glu (N-term), deamidated (NQ), and acetyl (Protein N-term). Two missed tryptic cleavages were allowed, and the protein-level false discovery rate (FDR) was ≤ 1%.

### Histology and immunofluorescence staining

Hearts were fixed in 4% paraformaldehyde (PFA) at 4°C overnight. Tissues were then rinsed with 1X PBS and cryoprotected in 30% sucrose/1X PBS at 4°C overnight before embedding in OCT (Tissue-Tek, Cat. #4583). Five micron sections were collected and subjected to either picro sirius red staining or immunofluorescent staining.^3^ Images of picro sirius red staining were acquired using Olympus BX69 microscope with NIS Elements software, and fibrotic area in each image was determined using Image J (NIH free software). Images for immunofluorescent staining were acquired using an inverted Nikon A1R confocal microscope and quantified with NIS Elements AR 4.13 software.

### Transmission electron microscopy

Hearts of anesthetized mice were perfused with 1% paraformaldehyde/2% glutaraldehyde (vol/vol) in cardioplegic solution (50 mmol/L KCl, 5% dextrose in 1X PBS), followed with 1% paraformaldehyde/2% glutaraldehyde (vol/vol) in 0.1 mol/L cacodylate buffer, pH 7.2. The heart was then removed and scars were isolated, divided into small fragments and fixed in 1% paraformaldehyde/2% glutaraldehyde (vol/vol) in 0.1 mol/L cacodylate buffer, pH 7.2 at 4 °C, followed by post fixation in 2% OSO^4^ (in 0.1mol/L cacodylate buffer) before dehydration in acetone and embedding in epoxy resin. Ultrathin sections were counterstained with uranium and lead salts. Images were acquired on a Hitachi 7600 electron microscope equipped with an AMT digital camera.

### Atomic force microscopy

Hearts were fixed in 4% paraformaldehyde overnight, embedded in OCT (TissueTek), and cryosections of 30 microns were mounted and dried onto gelatin-coated glass coverslips. Sections were rinsed in 1X PBS to remove OCT prior to imaging. Data was obtained using a Bruker NanoWizard 4 XP AFM system integrated with a Nikon Eclipse Ti inverted microscope and analyzed using JPK Data Processing software (version 7.0.101). Tissue stiffness was determined using an MLCT-BIO-E probe with a spring constant in the range of 0.061-0.064 N/m in Force Mapping mode. A 5×5 micron area was probed at 64 different points using a set point of 1.2 nN, a z length of 1.5 mm and z speed of 1 µm/sec. At least 3 areas of left ventricle free wall myocardium were scanned per section, giving a total of 150-250 data points, and values were averaged for each mouse. Young’s Modulus (kPa) was calculated by fitting force curves to the Hertz model, with corrections for quadratic tip shape/size, offset in height, and tip approach angle using JPK Data Processing software.

### Myocyte contractility

Cardiomyocytes were isolated by Langendorff perfusion, as described previously.^22^ Briefly, mice were injected with 100 U heparin and 10 minutes later hearts were excised, mounted to a cannula and perfused with collagenase digestion buffer. Dissociated hearts were run through a 100-micron mesh filter and cardiomyocytes allowed to settle for 10-15 minutes, rinsed and calcium reintroduction performed prior to taking measurements. Using an IonOptix data acquisition system for sarcomere detection, myocytes were paced at 10V and 1Hz; at least 8 myocytes were recorded per heart, and traces were analyzed using Ion Wizard software (IonOptix, version 7.5.3.165).

### Force measurements on decellularized tissue strips

To examine force generation of myocardial ECM, the left ventricular free wall was cut into 3 mm (length) x 2 mm (width) strips using a slicing mold, dissecting scope and ruler; 5-6 strips were obtained per heart. The strips were rinsed in 1X PBS and then digested for ∼48h in 1% SDS/1X PBS containing cOmplete™ Mini protease inhibitor cocktail (Millipore Sigma, Cat.# 4693124001) and PhosSTOP phosphatase inhibitor cocktail (Roche Diagnostics, Cat.# 4906837001). Since decellularization caused strips to lose some of their shape, strips were re-measured and trimmed to obtain 3 mm (length) x 2 mm (width) x 1 mm (height) for the experiment, and at least 4 strips were measured per heart. The decellularized tissue strips were attached to aluminum t-clips (Kem-Mil #1870) and mounted onto a muscle fiber test apparatus (Aurora Scientific, Model 802D-160-322) such that initial tension of each strip was set to zero. Tissue length was increased 5% over 50 ms, held for 150 ms, then returned to baseline tension. This was repeated at 5% increments from 0 to 60% length increase for each strip. Force was monitored using DMC v600A software (Aurora Scientific). Change in force was calculated as the difference between max force generated after the 50 ms pull and the minimum force achieved after each relaxation period. Minimum force was determined when the rate of force decay was zero and relaxation rate was calculated by solving for the derivative of the best fit second-degree polynomial trend line. The slope of the derivative is the relaxation rate.

### Cardiac function by echocardiography and invasive hemodynamics

Mice were anesthetized with 1.5% isoflurane and subjected to 2D guided M-mode echocardiography using a VisualSonics Vevo 3100 Imaging System with a 40-MHz transducer as described.^6^ Data were obtained by personnel blinded to genotype and treatment.

For invasive hemodynamics mice were anesthetized with 2.5% isoflurane. A high fidelity, solid state 1.2F pressure volume (PV) catheter (Transonic Scisense Inc) was inserted into the left ventricle via right carotid cutdown and retrograde introduction of the PV catheter into the LV. The signal was optimized by phase and magnitude channels.^23^ Mice are normalized to 1.5% isoflurane and 37 °C for at least 5 minutes, at which point data were collected by PowerLab 8/36 (ADInstruments) and LabChart 7 Pro (ADInstruments). A minimum of 10 continuous seconds of recorded data were averaged for each time point.

### Statistical analyses

Data are expressed as mean ± SEM unless otherwise stated. mRNA and protein expression levels were normalized to GAPDH, unless otherwise stated. The following tests were performed using Graphpad (Prism 9). Student’s t-test was performed for two group comparisons. One-way ANOVA with Tukey’s post hoc analysis was performed for multiple group comparisons as well as to determine the adjusted P value between group comparisons. Statistical significance for each experiment is described in figure legends.

## Results

### *Col1a2*^*-/-*^ mice develop cardiomyopathy and a compromised cardiac ECM

To investigate the function of type I collagen in the heart, we generated a mouse line carrying a *Col1a2* “knockout first allele”, see methods (Fig. 1A). The knockout first construction typically results in a null allele due to the insertion of a large coding cassette between the first coding exon and the rest of the gene. Indeed, RNA from whole ventricles of wild-type (*WT*) and *Col1a2*^*-/-*^ mice showed a greater than 80% reduction of *Col1a2* expression by qRT-PCR (Fig. 1B). We also examined the impact of the loss of *Col1a2* transcript on total type I collagen in the adult heart at baseline. There was no difference in total type I collagen observed in frozen heart sections from *WT* versus *Col1a2*^*-/-*^ mice immunostained with an antibody against total type I collagen (Fig. 1C and D). This result is in agreement with previous findings whereby loss of collagen1a2 in mice is compensated by incorporation of an additional collagen 1a1 chain as an obligate trimer. ^10,11^

**Figure 1.**
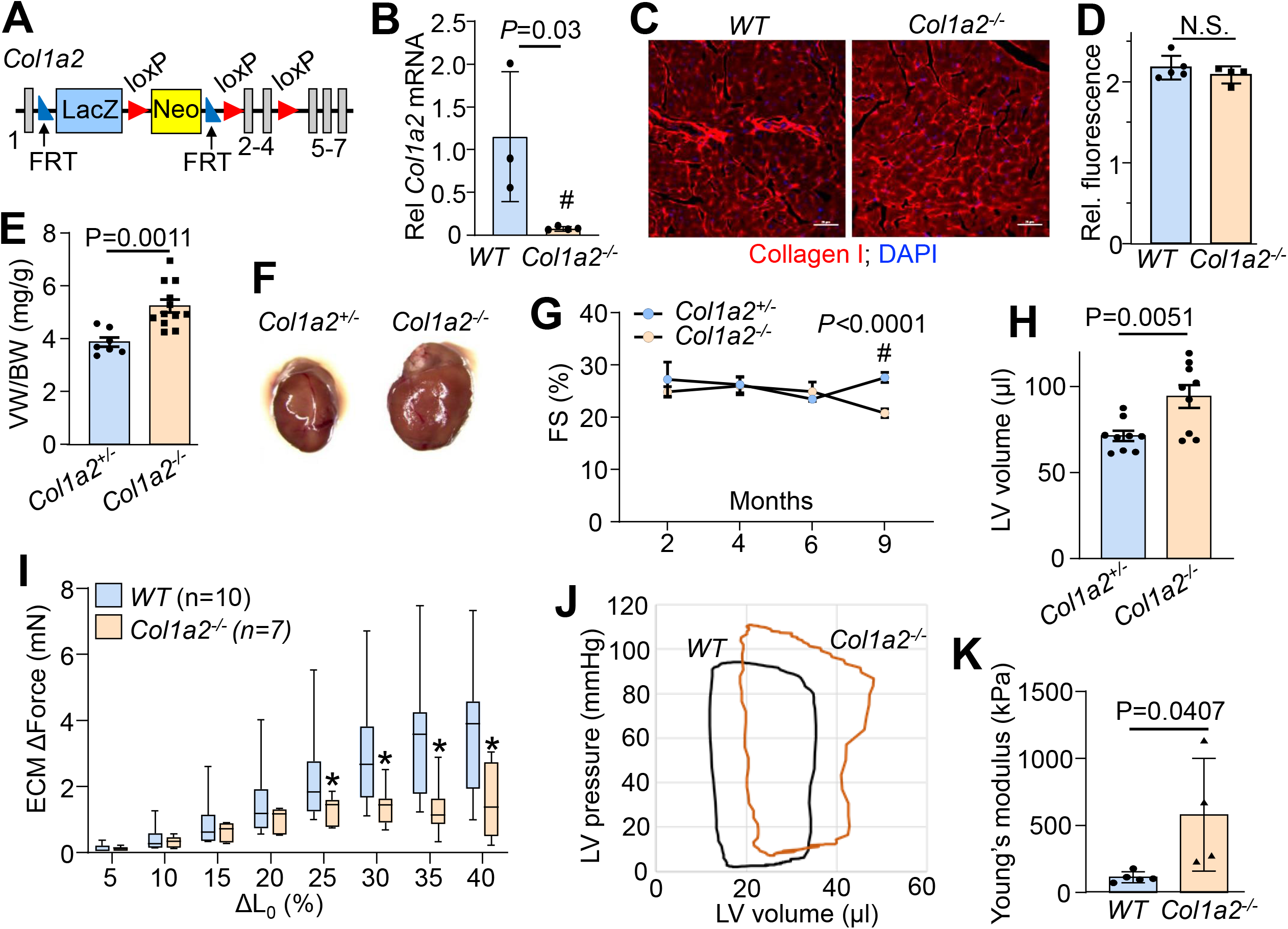
*Col1a2*^*-/-*^ mice develop cardiomyopathy and compromised cardiac ECM. (A) Schematic representation of the targeted *Col1a2* allele in the mouse that was used for both constitutive and tissue specific deletion of this gene. (B) Relative *Col1a2* mRNA expression in both wild-type (*WT*) and null (*Col1a2*^*-/-*^) mouse hearts at 3 months of age. (C) Representative immunohistochemistry from heart histological sections and quantification (D) of type I collagen in *WT* and *Col1a2*^*-/-*^ hearts at 3 months of age. Scale bar: 25μm. (E) Ventricle weight to body weight (VW/BW) ratio in *Col1a2*^*+/-*^ and *Col1a2*^*-/-*^ at 3 months of age. (F) Representative images of *Col1a2*^*+/-*^ and *Col1a2*^*-/-*^ mouse hearts at 3 months of age. (G) Echocardiographic measurement of cardiac fractional shortening (FS%) in *Col1a2*^*+/-*^ and *Col1a2*^*-/-*^ mice at 2, 4, 6 and 9 months of age, n=9 of each group at 9 months of age. (H) Echocardiographic measurement of left ventricular (LV) volume at diastole in *Col1a2*^*+/-*^ and *Col1a2*^*-/-*^ mice at 9 months of age, n=9. (I) Change in stretch-induced passive force generation of decellularized left ventricle strips (ECM) from *WT* and *Col1a2*^*-/-*^ mice at 8-10 weeks of age. One-way ANOVA analysis *P*=0.0032, \**P*<0.05, *Col1a2*^*-/-*^ versus *WT* with Tukey’s post-hoc analysis. (J) Representative baseline cardiac pressure-volume loops obtained by invasive hemodynamic measurements in hearts of *WT* and *Col1a2*^*-/-*^ mice at 8-10 weeks of age. (K) Using atomic force microscopy on cryosections, Young’s Modulus was determined for *WT* and *Col1a2*^*-/-*^ mouse hearts at 3 months of age. kPa, kilopascal. Student *t*-test for panels (B), (D), (E), (G) (H) and (K).

While both *Col1a2*^*+/-*^ and *Col1a2*^*-/-*^ mice are viable and grossly normal, *Col1a2*^*-/-*^ mice uniquely developed progressive cardiac hypertrophy by 10-12 weeks of age. The ventricle weight-to-body weight ratio increased more than 25% by 3 months of age compared to their *Col1a2*^*+/-*^ littermates (Fig. 1E). The hearts of *Col1a2*^*-/-*^ mice are also noticeably larger than *Col1a2*^*+/-*^ littermates on a morphological level (Fig. 1F). To examine the impact of the loss of *Col1a2*^*-/-*^ at a cellular level, we measured contractility of isolated cardiomyocytes and observed a significant reduction in fractional shortening at 8-weeks and 9-months of age (Supplemental Fig. 1A and B). This corresponded with a significant increase in cardiomyocyte thickness, but not length (Supplemental Fig1C and D). The reduction in contractility was also observed within the whole heart by echocardiography at 9 months of age (Fig 1G), along with dilation of the left ventricular chamber (Fig 1H).

To examine how loss of collagen 1a2 might compromise the structural rigor of the ECM in the mouse heart we measured stretch-induced passive force generated with decellularized left ventricle strips from *WT* and *Col1a2*^*-/-*^ mice (Fig. 1I). We observed a significant decrease in the passive force generated from *Col1a2*^*-/-*^ versus *WT* ventricle strips at increased lengths of stretching (Fig. 1I). This is again in agreement with previous findings that the left ventricle of OIM mice was more compliant and that they more readily expand with passive inflation of a balloon.^12^ With a more compliant cardiac ECM network, it was reasonable to hypothesize that the hearts in *Col1a2*^*-/-*^ mice would have shifted pressure-volume (PV) loop relationship. Indeed, using invasive hemodynamics we observed an upwards and rightwards shift in the PV-loops obtained from hearts of *Col1a2*^*-/-*^ mice, indicating increased afterload pressure with greater left ventricular distention. This phenotype was consistent with greater ventricular stiffness measured by atomic force microscopy (AFM), which showed a 4-fold increase in Young’s modulus in *Col1a2*^*-/-*^ hearts (Fig. 1K). Since the AFM measurements were performed with intact ventricular tissue, it suggests that cardiomyocytes compensate for the loss of ECM structural rigor, which secondarily leads to reduced myocyte and whole ventricular performance as observed in heart failure.^24^

### Progressive fibrotic response in *Col1a2*^*-/-*^ mouse hearts

Given the observed progressive cardiomyopathy observed in *Col1a2*^*-/-*^ mice we also interrogated the underlying molecular basis of this response by changes in gene expression with mRNA microarrays. Interestingly, the most prominent changes were in genes related to the fibrotic response or genes that impacted mechanical properties of the ECM. Cardiac ventricular gene expression analysis showed only 225 genes upregulated and 249 genes downregulated in the heart due to loss of *Col1a2* (Supplemental Table 1). Gene Ontology (GO) enrichment analysis showed increased expression of a group of structure-and ECM-altering genes that included *Acta1, Postn, Fbn1, Loxl, Loxl2, Mmp19, Timp1* and *Sparc* (Fig. 2A). Furthermore, we performed ECM mass spectrometry using whole left ventricles from hearts of either *Col1a2*^*+/-*^ or *Col1a2*^*-/-*^ mice. Consistent with the mRNA analysis, only a handful of ECM proteins were found to be increased in *Col1a2*^*-/-*^ groups (Fig. 2B). One of the more interesting genes was that of periostin, which typically is induced to mediate collagen maturation after injury.^25^ Indeed, periostin deposition was uniquely detected in *Col1a2*^*-/-*^ adult hearts by both immunohistochemistry and western blotting (Fig. 2C and D). Together these findings indicate that in the absence of collagen 1a2, the heart attempts to compensate for the structural weakness by inducing additional fibrotic genes or ECM maturation genes.

**Figure 2.**
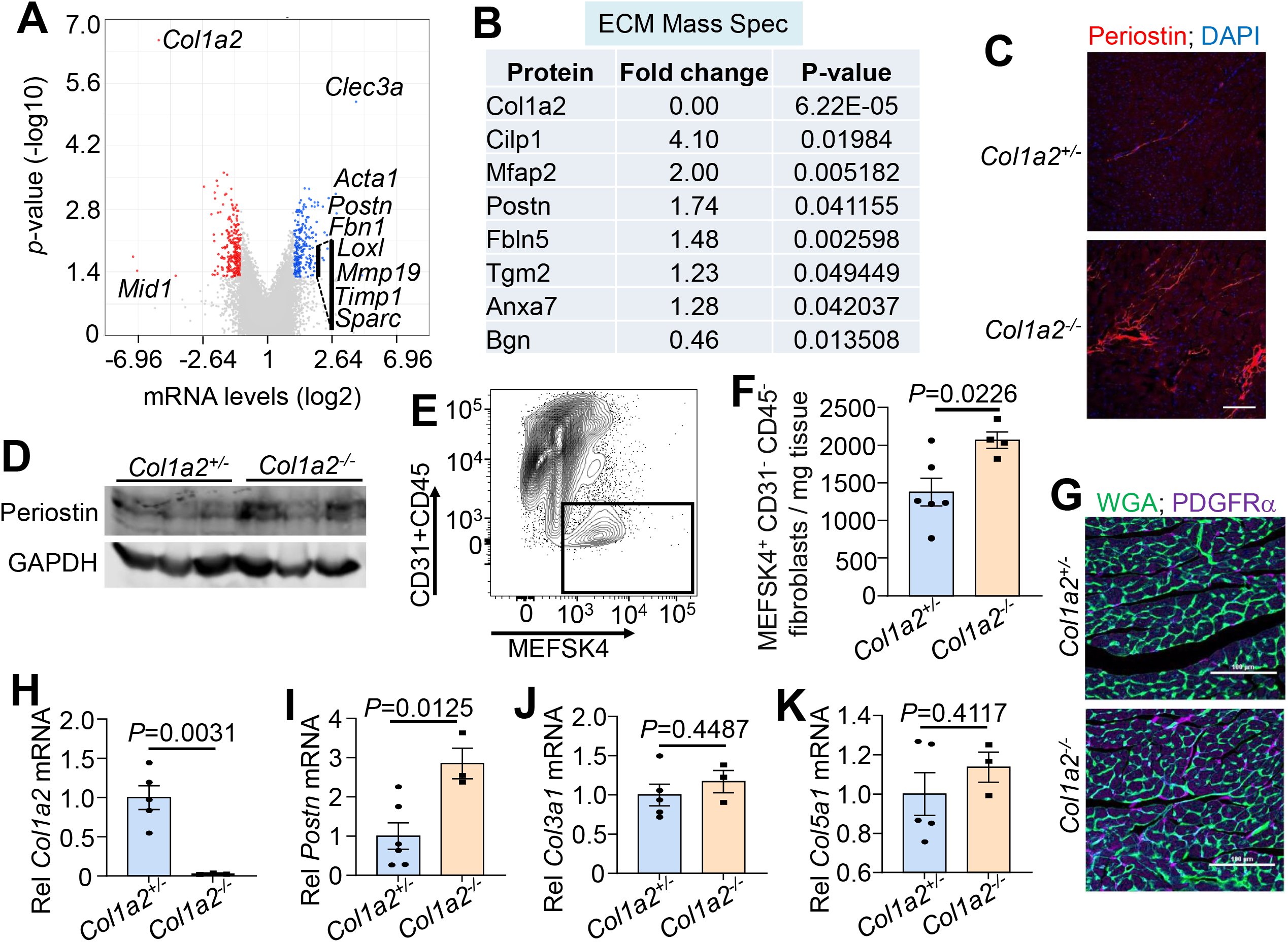
Activation of the fibrotic program in *Col1a2*^*-/-*^ mouse hearts. (A) Whole ventricle mRNA microarray analysis of *Col1a2*^*-/-*^mouse hearts compared to *Col1a2*^*+/-*^ at 2 months of age, n=3 per genotype. (B) Mass spectrometry analysis of ECM protein changes in *Col1a2*^*-/-*^ mouse hearts compared to *Col1a2*^*+/-*^ hearts at 3 months of age, n=4 per genotype. (C) Representative immunofluorescence images and (D) Western blot analysis of periostin from hearts of *Col1a2*^*+/-*^ and *Col1a2*^*-/-*^ mice at 3 months of age. Scale bar: 25 µm. (E) Flow cytometric gate strategy and (F) analysis of cardiac fibroblasts (MEFSK4^+^/CD31^-^/CD45^-^) from dissociated hearts of *Col1a2*^*+/-*^ and *Col1a2*^*-/-*^ mice at 3 months of age. (G) Representative immunofluorescence images of platelet-derived growth factor receptor (PDGFR)-α (purple) in *Col1a2*^*+/-*^ and *Col1a2*^*-/-*^ mice at 3 months of age. Wheat germ agglutinin (WGA) staining is green and shows outlines of cardiomyocytes. Scale bar: 100 µm. Relative mRNA expression of *Col1a2* (H), *Postn* (I), *Col3a1* (J) and *Col5a1* (K) in sorted cardiac fibroblasts (MEFSK4^+^/CD31^-^/CD45^-^) from *Col1a2*^*+/-*^ and *Col1a2*^*-/-*^ mice at 9 months of age. Student *t*-test for panels (F), (H), (I), (J) and (K).

The increase in fibrosis in adult *Col1a2*^*-/-*^ hearts suggested activation of the cardiac fibroblast. We examined the number of fibroblasts in both *Col1a2*^*+/-*^ and *Col1a2*^*-/-*^ hearts at baseline using flow cytometry to quantify the number of CD31^-^/CD45^-^/MEFSK4^+^ cells (Fig. 2E). Cells were isolated, counted and normalized to tissue weight, showing a ∼30% increase within the left ventricles of *Col1a2*^*-/-*^ compared to *Col1a2*^*+/-*^ mice at 3 months of age (Fig. 2F). Heart sections from either *Col1a2*^*+/-*^ or *Col1a2*^*-/-*^ mice were also stained with a PDGFRα antibody, which is largely expressed by fibroblasts in the heart, and this analysis showed an obvious increase in staining in hearts from *Col1a2*^*-/-*^ mice (Fig. 2G). Indeed, while *Col1a2* mRNA was still absent at 9 months of age (Fig 1H), the gene expression signature of fibroblast activation for periostin (*Postn*) was also increased in *Col1a2*^*-/-*^ hearts compared with control *Col1a2*^*+/-*^ mice at baseline (Fig. 2I). However, activation of 2 other major collagen isoforms in the heart remained unchanged through 9 months of age (Figs. 2J and 2K). Collectively, these results suggest that in the absence of collagen 1a2 in the heart, fibroblasts are activated where they attempt to compensate for the reduction in ECM structural rigor, possibly through expression of genes detected in our microarray and Mass Spec analysis (Fig. 2A and B).

### Greater cardiac fibrosis with pressure overload stimulation

We also investigated how *Col1a2*^*-/-*^ mice respond to acute pressure overload injury, which is a known inducer of collagen production and fibrosis. Eight-week-old *Col1a2*^*+/-*^ and *Col1a2*^*-/-*^ mice underwent transverse aortic constriction (TAC) surgery to induce pressure overload hypertrophy, and the hearts were harvested one week later for further analysis (Fig. 3A). TAC injury induced a strong fibrotic response and significant cardiomyocyte hypertrophy in controls. Heart sections were stained with anti-type I collagen antibody (Fig. 3B), which showed a significant increase in *Col1a2*^*+/-*^ controls compared with no increase in deposition over 1 week of TAC in *Col1a2*^*-/-*^ hearts (Fig. 3C). These results suggest that during an acute injury, α1(I)-containing homotrimer collagen fibrils do not have sufficient time to compensate for the loss of α2(I) during the acute phase of an injury response. However, *Col1a2* and *Col1a1* are normally induced by pressure overload stimulation in WT hearts over 1 weeks’ time compared with a sham procedure, similar to *Postn, Col3a1* and *Col5a1*, yet this fibrotic induction profile was even greater in mice lacking the *Col1a2* gene (Fig 3E-H). The cardiac hypertrophic program was also further increased after TAC stimulation in *Col1a2*^*-/-*^ mice compared with *Col1a2*^*+/-*^ controls, although it was the same relative increase from the Sham controls of each genotype (Fig 3I). These results further support the conclusion that in the absence of structurally rigorous type I collagen in the heart, a secondary reactive fibrotic program is initiated, that secondarily leads to cardiomyopathy.

**Figure 3.**
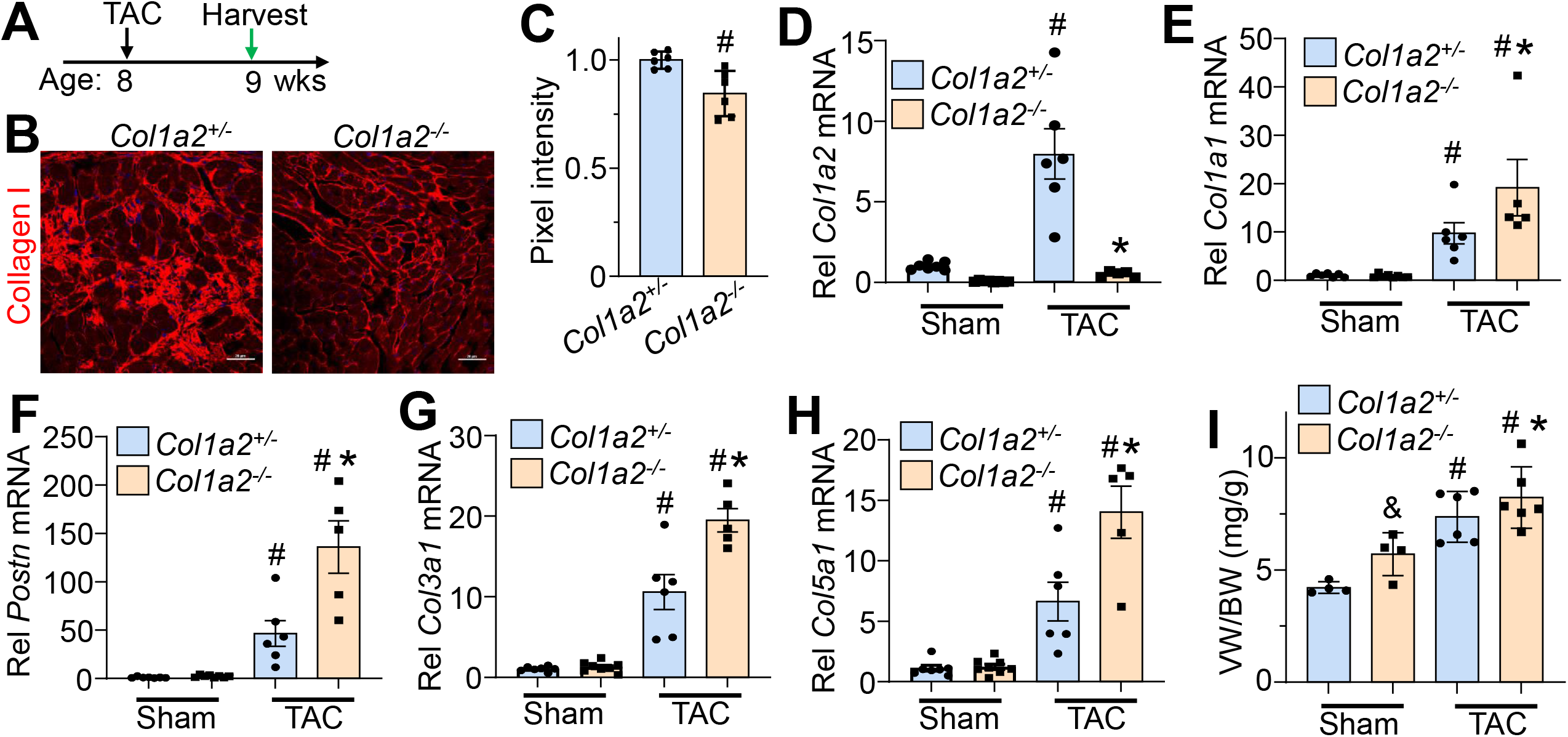
*Col1a2*^*-/-*^ mice show enhanced pressure overload hypertrophy. (A) Schematic representation of the experimental design in germline *Col1a2*^*-/-*^ mice with TAC surgery at 8 weeks of age and harvest at 9 weeks. (B) Representative immunohistochemistry images from cardiac histological sections with quantification (C) of type I collagen from hearts of *Col1a2*^*+/-*^ and *Col1a2*^*-/-*^ mice 1 week after TAC surgery. Scale bar: 20 µm. ^#^*P*<0.05. Relative mRNA expression of *Col1a2* (D), *Col1a1* (E), *Postn* (F), *Col3a1* (G) and *Col5a1* (H) from hearts of *Col1a2*^*+/-*^ and *Col1a2*^*-/-*^ mice at 9 weeks of age. (I) Measurements of ventricle weight to body weight ratio (VW/BW) in the indicated groups of mice. For panels (D) to (I), One-way ANOVA analysis *P*<0.05, with Turky’s post-hoc analysis for between groups comparison. ^#^*P*<0.05, compared to sham *Col1a2*^*+/-*^; \**P*<0.05, *Col1a2*^*+/-*^vs *Col1a2*^*-/-*^TAC; ^&^*P*<0.05 *Col1a2*^*+/-*^ vs *Col1a2*^*-/-*^ in the sham group.

### Fibroblast-specific *Col1a2* deletion shows autonomous effect on hypertrophy

To specifically delete the *Col1a2* gene only in cardiac fibroblasts (or myofibroblasts) we generated *Col1a2-loxP* targeted mice by “flipping” out the intervening cassette containing the β-galactosidase gene and neomyocin resistance gene using the FRT site (Fig 1A). These newly generated mice were left with loxP (fl) sites flanking exons 2-4 of the *Col1a2* gene, which when crossed with mice harboring the tamoxifen-inducible *Tcf21-MerCreMer (MCM)* allele, permitted deletion of the *Col1a2* gene in resident cardiac fibroblasts at baseline (Fig. 4A, *Tcf21*^*MCM/+*^; *Col1a2*^*fl/fl*^ mice). These mice were further bred with *Rosa26*^*NG-loxP-eGFP*^ mice, so that the expression of eGFP was induced by the selected Cre allele (Fig. 4A, *Tcf21*^*MCM/+*^; *Col1a2*^*fl/fl*^; *Rosa26*^*NG/+*^ mice). Eight-week-old mice were fed tamoxifen chow for 3 weeks to ablate *Col1a2* gene expression specifically in cardiac fibroblasts, after which hearts were harvested at the end of week 17 (Fig. 4B). Heart histological sections from these mice were examined using fluorescence microscopy, which showed specific eGFP tracing of adult fibroblasts after tamoxifen only with the *Tcf21*^*MCM*^ allele (Fig. 4C). Western blotting with a collagen 1a2-specific antibody showed a reduction in expression of this protein in the hearts of *Tcf21*^*MCM/+*^; *Col1a2*^*fl/fl*^; *Rosa26*^*NG/+*^ mice with tamoxifen compared with the same mice not given tamoxifen (Fig. 4D). This mild reduction likely reflects slow turnover of type I collagen in the heart, the half-life of which has been estimated at about 45 days in muscle.^26^ Interestingly, even this intermediate level of reduction of collagen had an impact on heart weight (Fig 4E). While there were no differences between control and tamoxifen-chow fed groups at 8 weeks of age, we observed that fibroblast-specific *Col1a2* deleted mice had a significantly lower ventricle weight to body weight (VW/BW) at 17 weeks of age (i.e., no differences in body weight). These findings suggest that a well-organized ECM is required for cardiomyocyte homeostasis in adulthood. However, we also hypothesize that given enough time, a reactive secondary phase would likely progress given our observations in germline *Col1a2*^*-/-*^ mice when present through development through 10-12 weeks of age (as shown in Fig 1).

**Figure 4.**
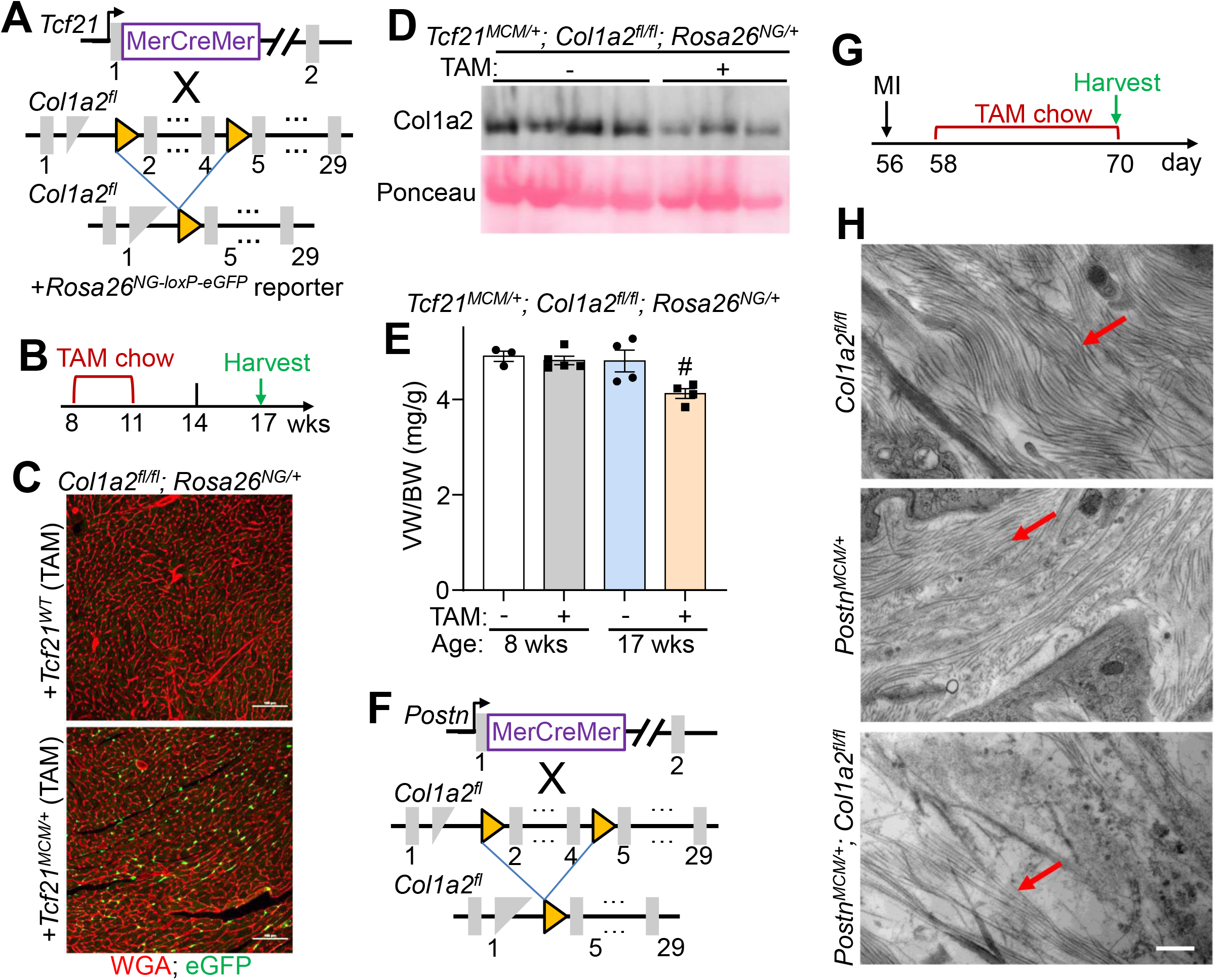
Fibroblast-specific and adult *Col1a2* deletion and heart analysis. (A) Schematic representation of the *Col1a2* loxP allele and resulting exon 2-4 deletion with a celltype specific Cre line, such as the *Tcf21*^*MCM*^ allele. These mice were also contain the Rosa26^NG-loxP-eGFP^ reporter. (B) Timeline for tamoxifen (TAM) chow feeding and harvesting of the experiment. (C) Immunofluorescent staining from cardiac histological sections for WGA (red) and eGFP from *Rosa26*^*NG-loxP-eGFP*^ reporter from the indicated groups of mice post-tamoxifen. Scale bar: 100 µm. (D) Western blotting from decellularized ECM for collagen 1a2 in hearts from the indicated groups of mice. (E) Measurements of ventricle weight to body weight ratio (VW/BW) at 8 weeks and 17 weeks of age in the indicated groups of mice. One-way ANOVA analysis *P*<0.05, with Turky’s post-hoc analysis ^#^*P*<0.05, mice with tamoxifen chow compared to normal chow at 17 weeks old. (F) Schematic of cell specific Cre induction to ablate *Col1a2* in activated cardiac fibroblasts (myofibroblasts) expressing the *Postn*^*MCM*^ *allele*. (G) Experimental timeline with MI (myocardial infarction) and TAM chow feeding followed by harvest. (H) Transmission electron microscope images of histological section from infarcted myocardium 2 weeks post-MI. Red arrows indicate collagen fibers within the scar region. Scale bar: 800 nm.

To more specifically examine fibrosis in the adult heart as mediated by activated cardiac fibroblasts, and how this might impact the integrity of type I collagen and cardiac hypertrophy, we used the tamoxifen-inducible *Postn-MerCreMer* gene inserted mice (*Postn*^*MCM*^) (Fig. 4F). Mice at 8 weeks of age were subjected to myocardial infarction (MI) injury and fed tamoxifen chow 2 days post-surgery until tissue harvesting 2 weeks later (Fig. 4G). Importantly, transmission electron microscopy of the MI scar region at 2 weeks after injury showed abundant organized collagen fibrils in *Postn*^*MCM/+*^ and *Col1a2*^*fl/fl*^ control mice, but not in *Postn*^*MCM/+*^; *Col1a2*^*fl/fl*^ mice (all mice are given tamoxifen) (Fig 1H). These results indicate that deletion of *Col1a2* from myofibroblasts impedes the production of new type I collagen fibrils in the MI scar region of the heart.

Since adult-specific deletion of *Col1a2* from fibroblasts of the heart resulted in defective collagen production, we again re-investigated the effects on cardiac hypertrophy with TAC stimulation. However, this time we attempted to address the temporal aspects of the effect between ECM structural properties and cardiomyocyte hypertrophy given the apparent contradictory finding of less hypertrophy in adult *Tcf21*^*MCM*^ mediated deletion of *Col1a2* (Fig 4E) versus greater hypertrophy with germline and early developmental deletion of *Col1a2* (Fig 1E). *Postn*^*MCM/+*^;*Col1a2*^*fl/fl*^ and control lines were subjected to TAC surgery at 8 weeks of age and mice were injected (i.p.) with tamoxifen followed by chow feeding up until sacrifice at 1, 2 or 6 weeks post-surgery (Fig. 5A). We observed strong upregulation of type I collagen immunostaining in the hearts of control *Col1a2*^*fl/fl*^ mice 1 week after TAC, while this upregulation was lacking in *Postn*^*MCM/+*^; *Col1a2*^*fl/fl*^ mice (Fig. 5B). While both groups showed a significant increase in VW/BW ratios 1 week post-TAC surgery, the increase was significantly reduced in the *Postn*^*MCM/+*^;*Col1a2*^*fl/fl*^ group (Fig. 5C). These results are consistent with the need for increased ECM content or rigor in the heart to support productive cardiac hypertrophy as observed previously.^5, 6^ Interestingly, two weeks of TAC stimulation showed a more intermediate profile where fibrosis was now only partially reduced in *Postn*^*MCM/+*^;*Col1a2*^*fl/fl*^ mouse hearts versus control *Col1a2*^*fl/fl*^ (Fig 5D and E). However, by 2 weeks there was no longer a significant difference in VW/BW ratios between *Col1a2*^*fl/fl*^ and *Postn*^*MCM/+*^;*Col1a2*^*fl/fl*^ groups (Fig. 5F), which was also the case 6-weeks post-TAC surgery (Fig. 5G). At 6 weeks post-TAC, total type I collagen staining remained decreased in *Postn*^*MCM/+*^;*Col1a2*^*fl/fl*^ mouse hearts (Fig. 5H), although total fibrosis was now fully compensated between the two groups (Fig. 5I). Collectively, these results suggest that acute inhibition of new ECM production with pressure overload stimulation attenuates cardiac hypertrophy, while continued fibrotic accumulation can compensate and eventually permits the same degree of cardiac hypertrophy over time.

**Figure 5.**
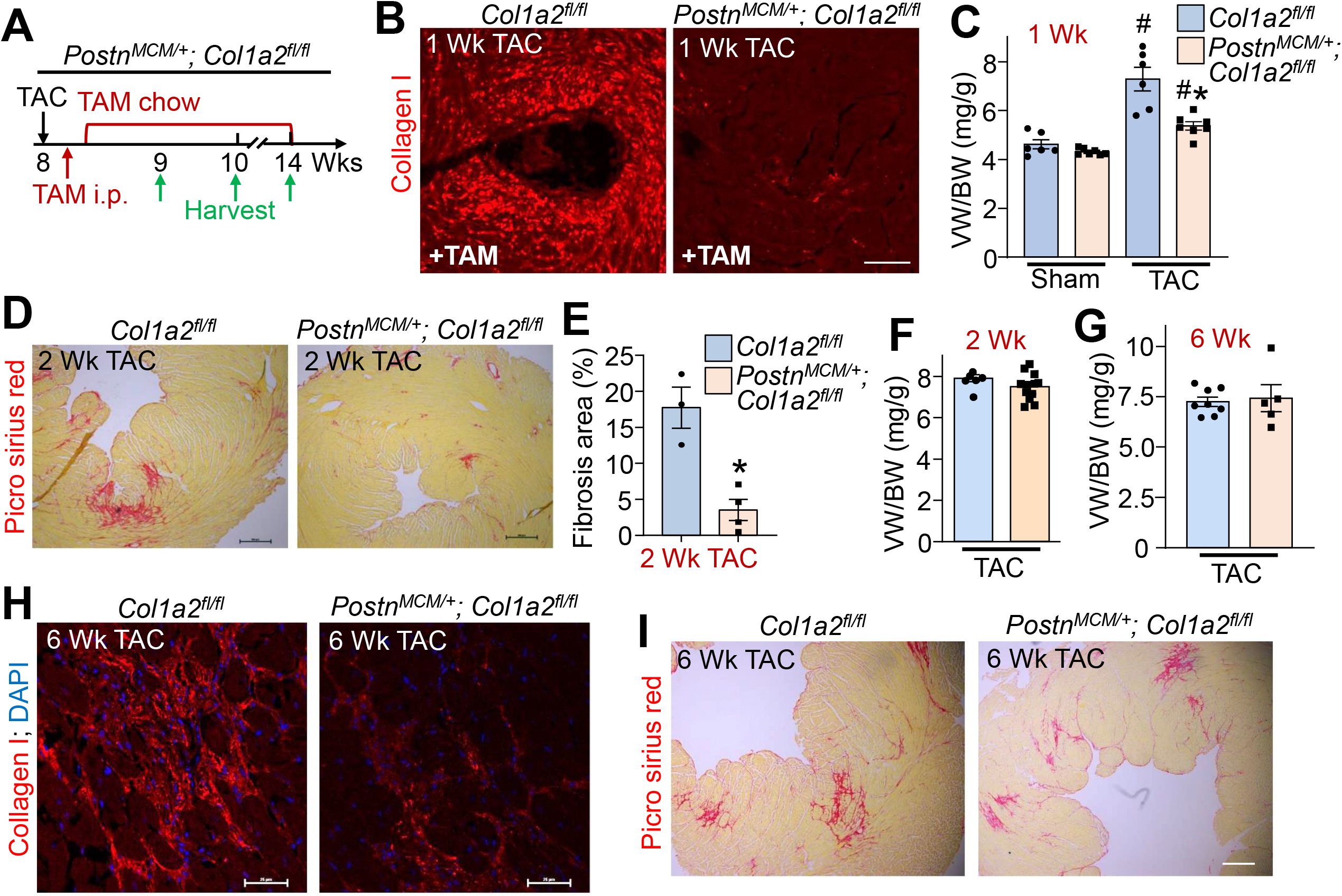
Myofibroblast-specific *Col1a2* deletion in the adult heart dynamically alters fibrosis and hypertrophy with pressure overload. (A) Timeline in *Postn*^*MCM/+*^*;Col1a2*^*fl/fl*^ mice at 8 weeks of age subjected to TAC, TAM administration/chow and tissue harvest. (B) Representative image of type I collagen staining (red) from heart histological sections 1 week post-TAC surgery. Scale bar: 25 µm. (C) Measurements of ventricle weight to body weight ratio (VW/BW) 1 week post-TAC surgery in *Col1a2*^*fl/fl*^ and *Postn*^*MCM/+*^*;Col1a2*^*fl/fl*^ mice. One-way ANOVA analysis *P*<0.05, with Turky’s post-hoc analysis ^#^*P*<0.05, compared to sham group with the same genotype; \**P*<0.05, *Col1a2*^*fl/fl*^ vs *Postn*^*MCM/+*^*;Col1a2*^*fl/fl*^ one week post-TAC surgery. (D) Picrosirius red staining in heart histological sections from *Col1a2*^*fl/fl*^ and *Postn*^*MCM/+*^*;Col1a2*^*fl/fl*^ mice 2 weeks post-TAC. Scale bar: 100 µm. (E) Quantification of Picrosirius red staining (fibrosis area %) in *Col1a2*^*fl/fl*^ vs *Postn*^*MCM/+*^*;Col1a2*^*fl/fl*^ mice 2 weeks post-TAC surgery; Student *t*-test, \**P*<0.05. Measurements of ventricle weight to body weight ratio (VW/BW) two weeks (F) and six weeks (G) post-TAC surgery in *Col1a2*^*fl/fl*^ and *Postn*^*MCM/+*^*;Col1a2*^*fl/fl*^ mice. (H) Representative images of type I collagen staining (red) from heart histological sections 6 weeks post-TAC surgery. Scale bar: 25 µm. (I) Picrosirius red staining in heart sections 6 weeks post-TAC. Scale bar: 100 µm.

## Discussion

Here we used a mouse model that altered collagen deposition and function in the heart to directly interrogate how this might impact cardiomyocyte growth dynamics. Unexpectedly, *Col1a2*^*-/-*^ mice showed a reactive expansion of the cardiac fibroblasts and secondary fibrosis in the heart due to a developmental absence of a structurally rigorous type I collagen-containing ECM network. However, acute deletion of *Col1a2* from the adult heart, specifically within fibroblasts with the *Tcf21*^*MCM*^ allele, reduced baseline heart weight normalized to body weight, suggesting that the structural properties of the ECM is sensed by cardiomyocytes even at baseline. Moreover, acute deletion of *Col1a2* from myofibroblasts in the adult heart with the *Postn*^*MCM*^ allele also produced less heart growth with 1 week of pressure overload, which was presumably due to the inhibition of new ECM production that normally occurs during this process.^2^ Indeed, we previously showed that inhibition of fibroblast activation and their ability to generate new ECM in the heart reduced hypertrophic growth ^3-6^. However, by 6 weeks of pressure overload the *Col1a2*^*-/-*^ mice generated enough additional collagen, albeit defective, to restore hypertrophic potential. Hence the quality and quantity of collagen within the heart are directly sensed by both cardiac fibroblasts and cardiomyocytes. This sensing by fibroblasts results in the enhanced fibrotic response and eventual cardiomyopathy observed with the developmental deletion of *Col1a2* as they age, while cardiomyocytes sense this same extracellular environment as they dynamically alter their size and growth properties.

A remaining question is how exactly the ECM influences various cell populations present within the myocardium during cardiac developmental growth or disease-based remodeling. During postnatal development of the heart, cardiomyocyte growth is matched with transient cardiac fibroblast activation that reciprocally drives ECM production in a process that allows the heart to mature and generate more force.^27^ Hence, it is likely that the cardiomyocyte directly senses the structural capacity of the ECM as it grows,^28^ which explains why acute inhibition of new ECM production results in less cardiac hypertrophy. In addition to ECM stiffness and structural support providing cardiomyocyte feedback, the ECM environment serves as a complex scaffold for latent growth factors that are released and activated during ECM remodeling, inflammation, or injury.^29, 30^ Transforming growth factor-β (TGFβ) is one such latent factor residing in the ECM, which when released directly programs fibroblast transformation to the myofibroblast, resulting in expansion of the ECM during disease stimulation.^31^ The epidermal growth factor family of latent growth factors are also released from the ECM where they can directly act on cardiomyocytes to program their hypertrophy, or by expansion of fibroblasts.^32^ Thus, our working hypothesis is that germline *Col1a2* null mice develop secondary hypertrophy, fibrosis and cardiomyopathy as they age due to a defect in the ECM that would initiate both potential mechanisms for altered growth and remodeling.

A properly organized collagen network binds fibrillin and the latent TGFβ binding proteins (LTBPs) that together help ensure the proper latency and release of TGFβ.^31^ Defective collagen would disrupt these complexes, likely leading to constitutive TGFβ release and ensuing fibrosis and disease, as well as release of other latent growth factors. This defect in the collagen network structural rigor would also likely be directly sensed by cardiomyocytes through their integrin and dystroglyan-sarcoglyan attachment complexes in stimulating internal signal transduction that could alter growth dynamics. Both mechanisms of affecting cardiac growth and remodeling are also likely in play during acute disease stimulation, as with 1 week of pressure overload. In this later case it appears that without new ECM production, myocytes are attenuated in their ability to fully hypertrophy, either due to less ECM structural rigor or less growth factor production and release from the ECM. In summary, the approach used here with constitutive and inducible *Col1a2* deletion demonstrated that a properly formed extracellular collagen network is critical for cardiac homeostasis and disease responsiveness to permit optimal compensation.

## Acknowledgements

None

---

**Supplemental Figure 1. Cardiomyocytes from *Col1a2***^***-/-***^ **mice are hypertrophic and have impaired contractility**. (A) Cell contractility was assessed in Langendorff-isolated individual cardiomyocytes from hearts of *Col1a2*^*+/-*^ vs *Col1a2*^*-/-*^ mice at 8 weeks and (B) 9 months of age. (C) Isolated cardiomyocyte length and (D) width measurements from *Col1a2*^*+/-*^ and *Col1a2*^*-/-*^ mice at 9 months of age. P values are as indicated, but the results in panel C are not significantly different. For panels (A) to (D), Student *t*-test.

