## Supplementary figures and images for "Cardiac fibroblasts regulate cardiomyocyte hypertrophy through dynamic regulation of type I collagen"

### Supplemental Figure 1

**Supplemental Figure 1**

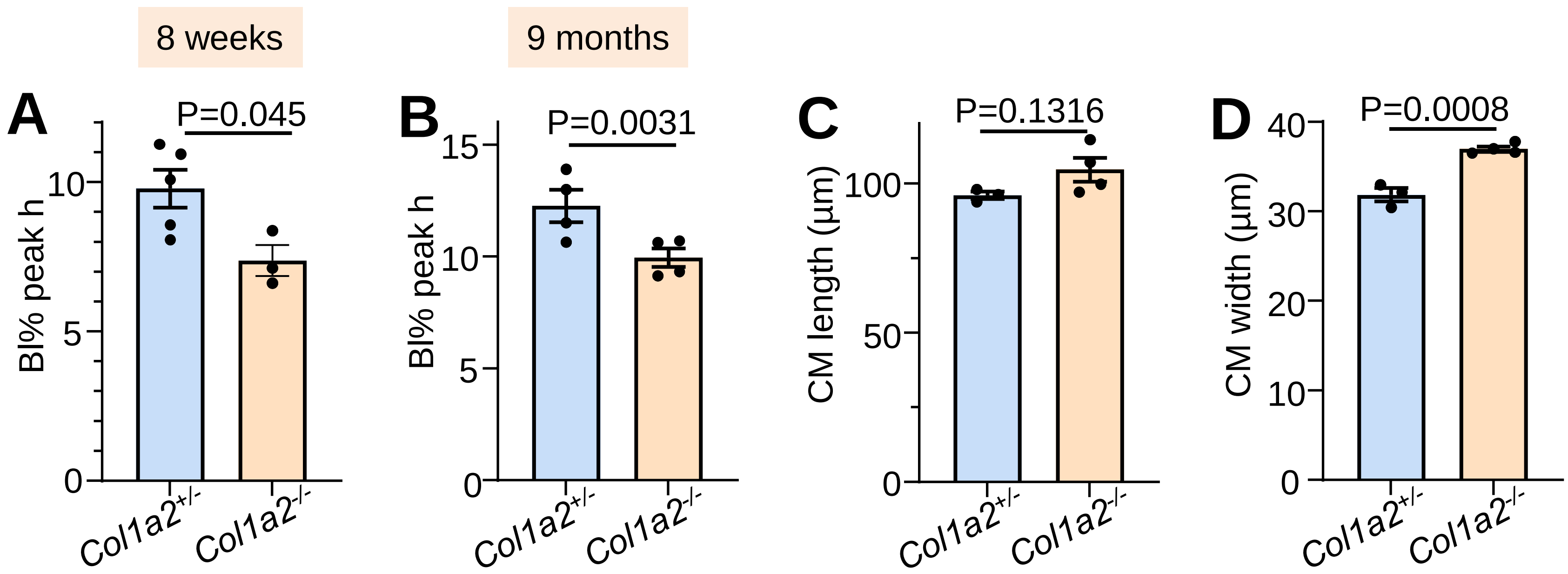
